# Fast but not furious: a streamlined selection method for genome edited cells

**DOI:** 10.1101/2021.01.22.427783

**Authors:** Haribaskar Ramachandran, Soraia Martins, Zacharias Kontarakis, Jean Krutmann, Andrea Rossi

## Abstract

In the last decade, Transcription Activator-Like Effector Nucleases (TALEN) and Clustered Regularly Interspaced Short Palindromic Repeats (CRISPR) based genome engineering have revolutionized our approach to biology. Due to their high efficiency and ease of use, the development of custom knock-out and knock-in animal or cell models is now within reach for almost every laboratory. Nonetheless, the generation of genetically modified cells often requires a selection step, usually achieved by antibiotics or fluorescent markers. The choice of the selection marker is based on the available laboratory resources, such as cell types, and parameters like time and cost should also be taken into consideration. Here, we present a new and fast strategy called **MAGECS** (magnetic-activated genome edited cell sorting), to select genetically modified cells based on the ability to magnetically sort surface antigens (i.e. tCD19) present in Cas9 positive cells. By using MAGECS, we successfully generated and isolated genetically modified human induced pluripotent stem cells (hiPSCs), primary human fibroblasts, SH-SY5Y neuroblast-like cells, HaCaT and HEK293T cells.

Our strategy expands the genome editing toolbox by offering a fast, cheap, and an easy to use alternative to the available selection methods.

## Introduction

Genome editing technologies have substantially improved the ability to make precise changes in the genomes of eukaryotic cells. Programmable nucleases, such as meganucleases derived from microbial mobile genetic elements, Zinc Finger (ZF) (Urnov et al. 2005; Miller et al. 2007) and Transcription activator-like effector nucleases (TALENs) (Boch et al. 2009; Moscou and Bogdanove 2009; Christian et al. 2010; Miller et al. 2011), have been used with discrete success to modify the genome of different species. In 2013, the CRISPR/Cas9 system from the type II bacterial adaptive immune system CRISPR system revolutionized our ability to interrogate the function of the genome (Cong et al., 2013; Mali et al., 2013). It was shown to be potentially useful clinically to correct genetic DNA mutations to treat diseases that are refractory to traditional therapies or where therapies are not yet available (Pires et al. 2016; Ma et al. 2017; Min et al. 2019).

The overall success of the CRISPR/Cas9 system compared to the other genome editing technologies lies in its overall efficiency, low cost, straightforward plasmid assembly and an unmatched number of available DNA targets (Cong et al. 2013). The main limiting factor for many labs, when using the CRISPR/Cas system, is the ability to sort transfected cells that carry the desired mutation.

Nowadays the enrichment of genetically modified cells after transfection is mainly achieved by antibiotic or fluorescence activated cell sorting (FACS). Both selection strategies are routinely used because they are relatively easy to perform and offer reproducible results. There are, however, several limitations that must be taken into consideration when choosing these selection methods. The most important ones are time, costs, and sorting efficacy. In order to simplify the selection process, we thought to exploit the use of a magnetic activated cell sorting system (Duda et al. 2014; Martin-Fernandez et al. 2020) to sort CRISPR genetically modified cells. Here, we describe a new method called “Magnetic Activated Genome Edited Cells Sorting” assay or MAGECS.

We show that MAGECS is a fast, easy and relatively inexpensive pipeline to select genetically modified cells, which can be used to enrich Cas9 positive cells in different cell types including hiPSCs, primary human fibroblast, HaCaT, neuroblast-like cells and HEK293T cells.

## Results

### Selection of genome edited cells

In order to select genome-engineered cells using MAGECS, we fused a truncated CD19 domain to the C-terminus of a pSpCas9-2A plasmid (Supplemental Fig. S1). Transfected cells expressed a functional Cas9, able to translocate into the nucleus (Fig. 1A) to mediate genome editing in the presence of a guide RNA (gRNA), as well as tCD19 surface marker, to enable the sorting of transfected cells. We tested MAGECS by transfecting HEK293T (Fig. 1B) and hiPSC cells (Fig. 1C) with the pSPCas9-2A-tCD19 plasmid. Immunostaining of transfected cells with CD19 antibody revealed the localisation of tCD19 at the plasma membrane. After magnetic sorting, the majority of eluted cells were CD19 positive cells (Fig. 1B-C) indicating the efficiency and purity of CD19 selection, with an average of 2 to 4 times enrichment, when compared to unsorted cells, depending on the cell type (Supplemental Fig S2).

**Figure 1.**
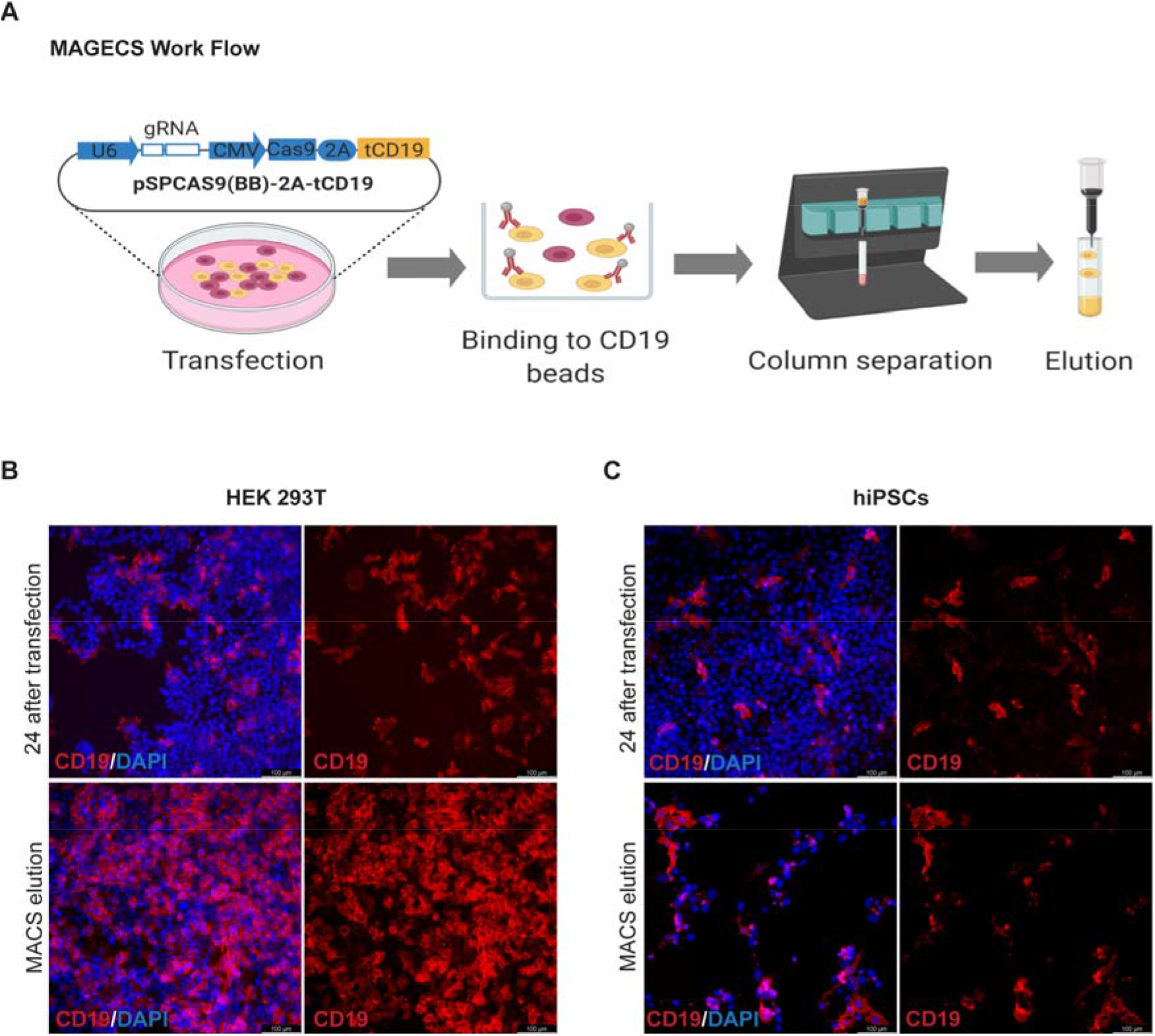
tCD19 cell surface localization after transfection ensures efficient sorting. **(A)** MAGECS work flow starting from transfection to elution. **(B-C)** Immunofluorescence of transfected HEK 293T cells. **(B)** and hiPSCs **(C)** using CD19 antibody (red) before and after MACS elution and nuclei (blue). Cell elute with CD19 signal indicates the specificity of the whole process.

### Magnetic sorting of genome edited HEK293T cells

Next, we designed and cloned different gRNAs targeting four human genes, *UDP Glucuronosyltransferase Family 1 Member A1 (UGT1A1)*, *dystrophin (DMD)*, *actin beta* (*ACTB)* and *centrosomal P4.1-associated protein (CPAP)* into the MAGECS plasmid vector (Fig. 2A). The MAGECS plasmids containing these gRNAs, were then transfected into HEK293T cells and the cells were MACS sorted after 48 hrs.

**Figure 2.**
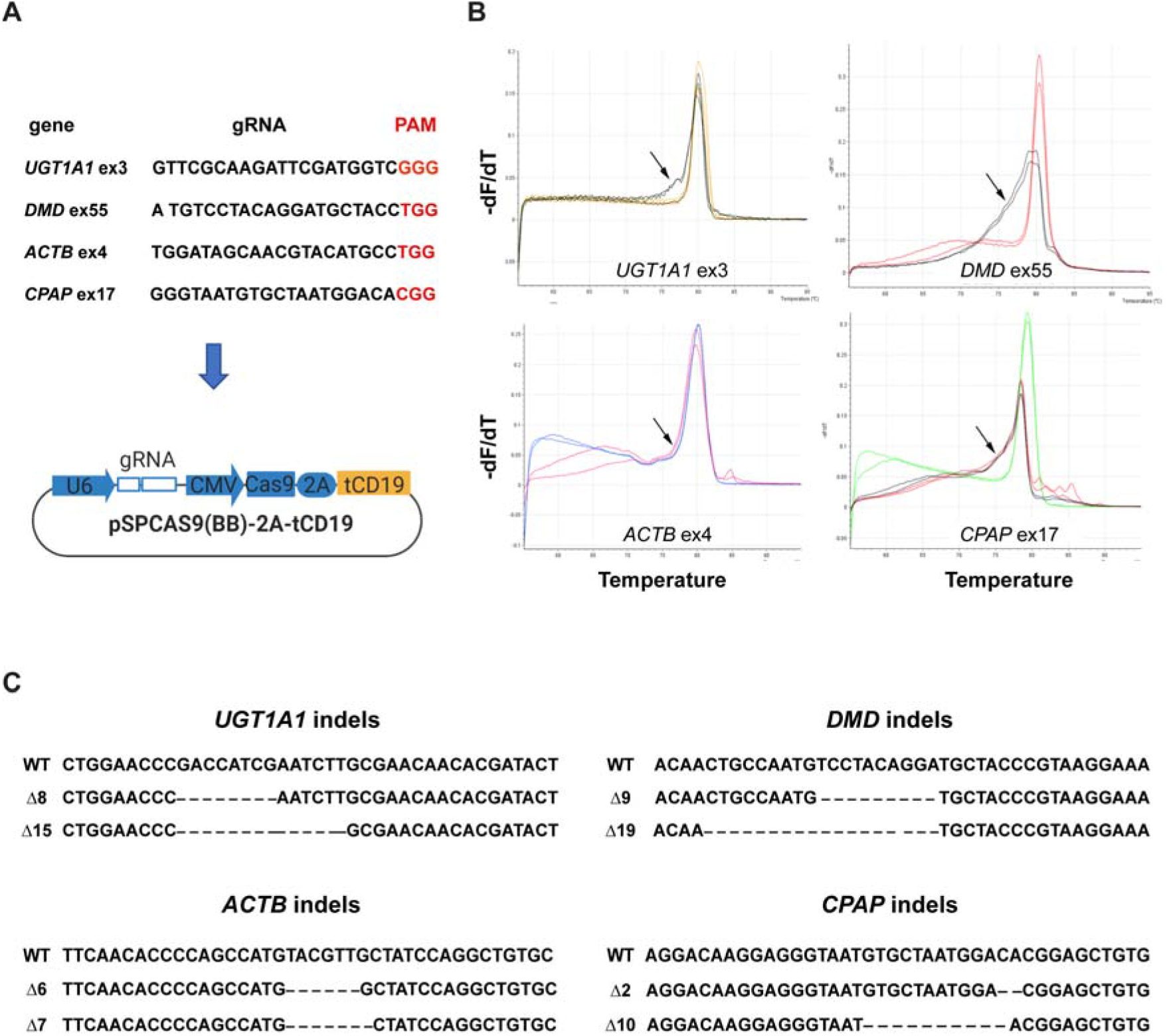
Efficient sorting of genome edited HEK 293T cells by MAGECS. **(A)** gRNA sequences targeting human *UGT1A1*, *DMD*, *ACTB* and *CPAP*. **(B)** HRMA curves showing the resolution of the genotypes. Arrows point to irregular HRMA curves which suggest the presence of DNA mutations. **(C)** Sanger sequencing of mutant HEK 293T cells.

In order to assess the genome editing efficiency, we took advantage of the High-Resolution Melting curve Analysis (or HMRA), which is routinely used because of its sensitivity, to detect known polymorphisms and particularly suitable for detecting indels induced by the genome editing technologies such as CRISPR/Cas9 system (Rossi et al. 2015).

After analysing wild-type (WT) and CD19 positive cells with HRMA a clear single pick was detected for WT and flow through (FT) cells indicating the lack of indels in absence of CD19 positve cells (Fig. 2B. Supplemental Figure S3). In contrast, CD19 positive cells displayed the typical irregular curve indicating the presence of DNA base indels (Fig. 2B, Supplemental Figure S3). Next generation sequencing and Sanger sequencing of CD19 positive cells confirmed the successful generation of indels indicating the efficacy of each gRNA and MAGECS to enrich genome edited cells (Fig. 2C).

### Magnetic sorting of genome edited primary cell lines

We next asked whether MAGECS would also be suitable for cell types beyond HEK293T cells, i.e. in particular primary cells, that are known to be more resilient to transfection and more sensitive to other sorting methods such as antibiotics or FACS.

To this end, primary human skin fibroblasts and SH-SY5Y neuroblast-like cells were transfected with gRNAs targeting *DMD* and *NADH:Ubiquinone Oxidoreductase Core Subunit S1* (*NDUFS1)*, respectively (Fig. 3A,D). Cells were magnetically sorted two days after transfection and divided in two halves. One half was plated on a new dish (Fig. 3B,E) and the other one was used to extract genomic DNA for HRMA (Fig. 3C,E). Indels were detected in CD19 positive cells by HRMA and Sanger sequencing (Fig 3 C,F). Furthermore, MAGECS did not change the morphology of primary human fibroblast nor neuroblast-like cells as shown in Fig. 3B, E.

**Figure 3.**
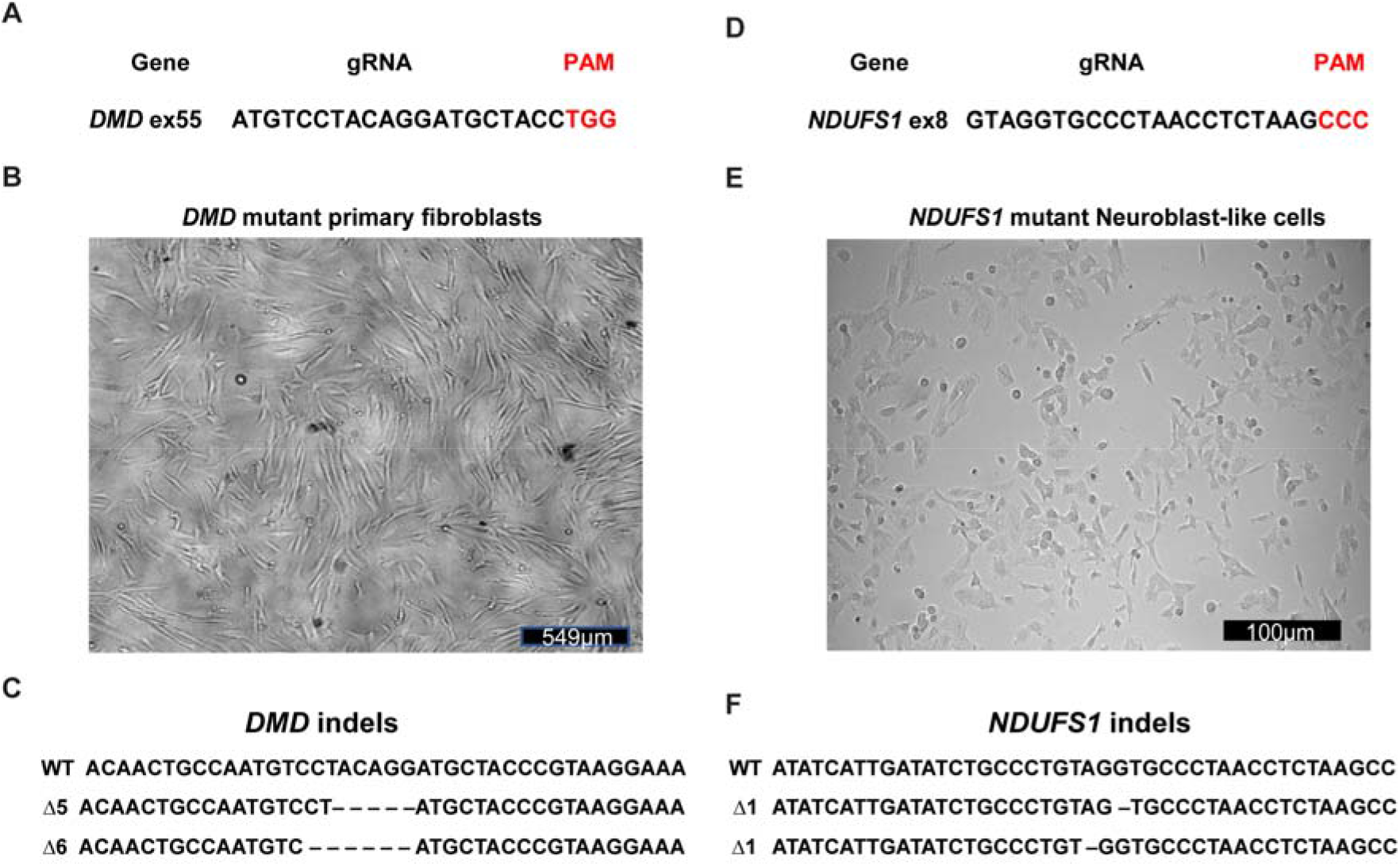
Efficient sorting of genome edited primary cells by MAGECS. **(A)** human *DMD* gRNA targeting sequence. **(B)** Brightfield micrographs of mutant primary fibroblasts after MAGECS. **(C)** Sanger sequencing of *DMD* mutant fibroblasts. **(D)** human *NDUFS1 g*RNA targeting sequence. **(F)** Brightfield micrographs of mutant neuroblast-like cells after MAGECS **(G)** Sanger sequencing of *NDUFS1* mutant neuroblast-like cells.

### MAGECS for sorting genome edited hiPSCs

hiPSCs are difficult to maintain in culture and to genetically modify their genome. Thus, we next asked whether MAGECS could be used to generate genome edited hiPSCs of single clonal origin with a defined mutation. Human iPSCs were transfected with two gRNAs targeting *UGT1A1* and *alpha fetoprotein* (*AFP*) (Fig. 4A) and magnetically sorted after 48 hrs. Single cells were plated in the presence of Rho Kinase (ROCK) inhibitors (i.e. Y-27632) onto a 96 multi-well plate and duplicated after two weeks (Fig. 4B). One plate was digested with Proteinase-K for further genotyping analysis and its duplicate was kept in the incubator. Deep sequencing analysis using an Illumina MiSeq sequencer (Schmid-Burgk et al. 2014) detected a high prevalence of mutations (data not shown) and revealed the presence of genetically modified cells carrying different types of mutations (Fig 4D). These data indicate that MAGECS can be used to easily and successfully genome engineer and sort hiPSCs. The viability and morphology of iPSCs was not affected by MAGECS (Fig. 4C) and further karyotype analysis of hiPSC clones indicated that MAGECS protocol did not result in any chromosomal abnormalities (data not shown).

**Figure 4.**
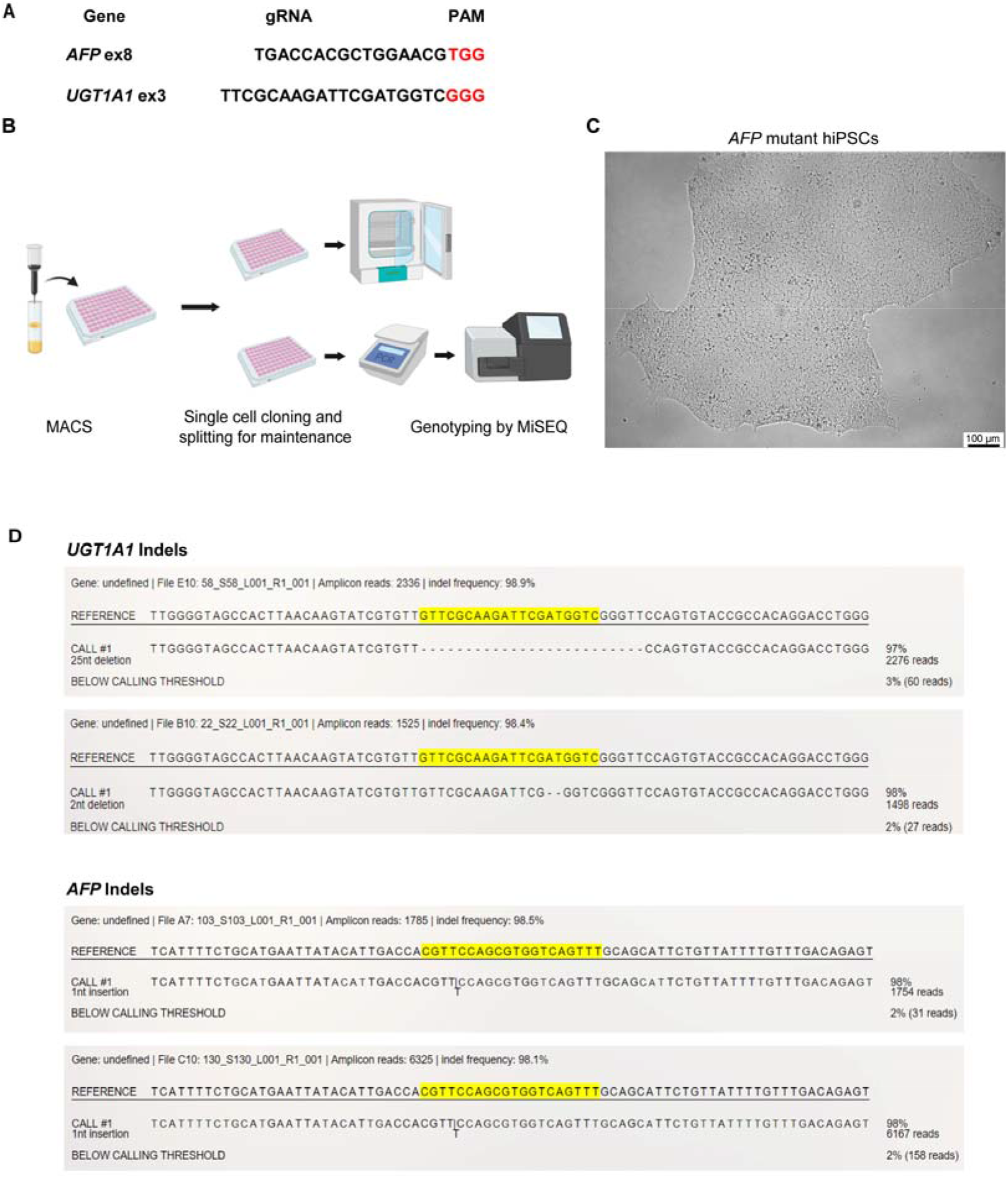
Efficient sorting of genome edited hiPSCs by MAGECS. **(A)** Human *UGT1A1* and *AFP* gRNA target sequences. **(B**) hiPSCs MAGECS pipeline representation for. **(C**) Brightfield micrographs of *AFP* mutant hiPSCs after MAGECS. **(D)** MiSEQ analysis of iPSCs *UGT1A1* and *AFP* mutants.

## Discussion

A major challenge when engineering mutant models, is the selection of genome edited cells or single clones carrying the desired mutation. By combining two already available tools, the CRISPR/Cas9 system (Mali et al. 2013; Doudna and Charpentier 2014; Cong and Zhang 2015) and magnetic sorting (Stefan Miltenyi 1990), we established a streamlined selection method of genome edited cells called MAGECS. Using MAGECS, we were able to enrich genome edited cells and select single clones for different cell types, including HEK293T, HaCaT, human primary fibroblasts and hiPSCs and gene targets. Colony formation was observed after single cell sorting by limited dilution, in all cell types tested, indicating that MAGECS does not affect the viability of cells. It also does not require expensive equipment or specialized personnel. The absence of genome edited cells in the flow through further supports efficiency and accuracy of this method in capturing tCD19 positive cells (Supplemental Fig. 2, Supplemental Fig. 3).

Our assay offers several advantages compared to the available selection methods. In comparison to antibiotics MAGECS is faster and more consistent. Antibiotics are affordable and can be used without specific equipment, but the cell selection requires time and it is laborious when compared with other selection methods (i.e. FACS or MACS). The first critical step, when using antibiotics, is to determine the optimal concentration to use for the selection of stable colonies. The optimal concentration can be determined after generation of a killing curve, so that un-transfected cells are efficiently cleared without affecting the transfected ones. Furthermore, antibiotics require a few days to clear cells that are not carrying the antibiotic resistant gene, making the process of generating genetically modified cells longer. Last but not least one of the main issues connected to the use of antibiotics is the possibility of spontaneously resistant clones that do not carry the gene of interest (Hawkey 1998).

Antibiotics-free selection using FACS can be an alternative to avoid the issues mentioned above. Accordingly, the use of FACS was shown to significantly increase the number of cells that can be screened in a short period of time. (Li et al. 2018). Unfortunately, the analysis of fluorescent markers requires large, complex, expensive instrumentation typically operated by highly trained specialists (Ren et al. 2019). The ability to quickly and simultaneously query multiple FACS parameters on large numbers of individual cells is generally reserved for shared core facilities or well-equipped specialized laboratories. Occasionally FACS sorted cells fail to form colonies due to the exposure to a strong laser beam and the high hydrostatic pressure (Smith et al. 2006). When compared with FACS, MAGECS has the great advantage to be very convenient from an economical point of view. We have estimated that the cost of a single MAGECS sorted sample is circa 6.5-7€ x 500k cells. In order to sort cells using MAGECS, a strong magnet (i.e. OctoMACS™ Separator), MACS columns and MACS microbeads are needed. The use of MAGECS allows us to sort up to 8 samples in parallel, making the whole procedure very fast and medium throughput. In this regard, another advantage offered by MAGECS is that it can be performed in the same facility where the cells are stored by virtually any member of the scientific staff since it requires only minimal training. Thus, not only reducing the costs, but also the stress inflicted on the cells.

FACS and MACS are both very robust methods and they are able to produce results that overall are very consistent (Sutermaster and Darling 2019). Nevertheless, the low cost of MAGECS and the ability to isolate genome edited cells with high specificity and by any staff member makes it an invaluable tool that will be useful for many scientists and labs working with genome editing.

In summary, we have demonstrated that MAGECS can sort genetically engineered cells with high efficiency and overall viability, and that it can be applied with minor protocol adjustments to a broad range of different cell types including hiPSCs and primary cells. We expect that the lower cell loss associated with MAGECS will represent an advantage which might make MAGECS particularly useful for those ‘stress sensitive’ hiPSCs reprogrammed from patients carrying certain diseases.

We thus believe that MAGECS will have the potential to become a widely used streamlined selection for the genome editing of cells.

## Methods

### Cell lines: maintenance and passaging

Human fibroblasts, HEK293T and HaCaT cells were maintained in Dulbecco’s Modified Eagle’s Medium (DMEM) (Gibco,) supplemented with 10% (v/v) heat inactivated fetal bovine serum (FBS) (Gibco,) and 1% (v/v) penicillin and streptomycin (P/S) (Pan Biotech, ref. P06-07100). SH-SY5Y neuroblast-like cells were maintained in Dulbecco's Modified Eagle Medium: Nutrient Mixture F-12 (DMEM/F-12) (Gibco) supplemented with 10% (v/v) FBS and 1% (v/v) P/S. The cells were passaged using 0.05% trypsin (Gibco). hiPSC line IMR90 (WiCell) was cultured in mTeSR medium (Stem Cell technologies) supplemented with 1% P/S on Matrigel-coated (Corning) plates. The medium was changed every day and the cells were passed every 5-6 days using ReLeSR (Stem Cell technologies). 24h prior transfection, iPSCs were passed using Accutase (Innovative Cell Technologies) in a well of a Matrigel-coated 6 well plate supplemented with 10μM Y27632 (Stem Cell technologies).

### Plasmids

pSPCas9-2A-puromycin plasmid was purchased from AddGene (ref 62988). The plasmid that we generated, called pSPCas9-2A-tCD19 or MAGECS plasmid, was obtained by using the pSPCas9-2A-puromycin and swapping the puromycin with tCD19. Briefly, pSPCas9-2A-puromycin plasmid was digested with EcoRI and gel purified without the puromycin cassette. tCD19 was then amplified by PCR using primers cold fusion adapters and cloned inside the gel purified pSPCas9-2A-puromycin backbone. The plasmid was then fully sequenced.

### Transfection

Cells were transfected in six-well plates (TPP, Switzerland) with 2μg of pSPCas9-2A-CD19 using FuGENE® HD Transfection Reagent (Promega) or Lipofectamine Stem Transfection Reagent (Thermo Fisher), in the case of iPSCs, for 48h according to the manufacturer’s instructions.

### MAC sorting

Cells were sorted with MACS using MS columns, OctoMACS separator and CD19 MicroBeads (human) (all from Miltenyi Biotec) modifying slightly the data sheet protocol according to the experimental needs and the cell types (figure 1).

After co-transfection, cells were detached (either with 0,05% trypsin or Accutase according to the cell type) and spun down at 500xg for 5min (HaCaT and HEK293T cells) or 300g for 10min (fibroblasts and hiPSCs). The pellet was then resuspended in 80μl MACS buffer (PBS pH 7.2, 0.5% FBS, and 2 mM EDTA) and 20μl CD19 Miltenyi MicroBeads, mixed well and incubated at 4°C for 15min. After the incubation, 1ml of MACS buffer was added to the cells and again spun down at 300xg for 10min. The supernatant was discarded and the pellet was resuspended in 500μl MACS buffer. After column equilibration, the cell suspension was loaded; CD19 negative cells passed through the column, while CD19 positive cells were retained. The retained cells were subsequently eluted using the syringe plunger to flush them trough. Cell suspension was spun down at 500g for 5 min (HaCaT and HEK293T) or 300xg for 10min (fibroblasts, SH-SY5Y and hiPSCs), re-plated into 6 well plates or μ-Slide 4 Well chambers (ibidi) in order to access the sorting efficiency.

### Cell lysis

The medium was discarded and the cells were lysed with the following of lysis buffer: 0.2 mg/ml proteinase K, 1mM CaCl_2_, 3mM MgCl_2_, 1mM EDTA, 1% Triton X-100, 10mM Tris (pH 7.5). Afterwards the reactions were first incubated for 10min at 55°C and then 15min at 95°C.

### Generation of CRISP/Cas9 mutants

gRNAs were designed using CRISPR design tool (https://www.crisprscan.org/) and cloned into the MAGECS plasmid. The resulting bicistrionic plasmid encoded the gRNA, the Cas9 nuclease and the surface tCD19 marker. gRNA activity and efficiency were assed suing High-Resolution Melt Analysis (Rossi et al. 2015) using a MyGo PRO real time PCR (IT-IS Life Science LTD).

### PCR Bar Coding

PCR was performed as previously described (Schmid-Burgk et al. 2014). Briefly, first-level PCR reactions were performed using 1μl PCR-compatible lysate as a template for a 6.25μl Q5 (NEB) PCR reaction according to the manufacturer’s protocol (annealing temperature: 60°C; elongation time: 15sec, 19 cycles) (NEB). Of this reaction, 2μl was transferred to a second-level PCR using the same cycling conditions and a combination of barcode primers that is unique for each clone. For all primer sequences see Supplemental Table S1.

### Deep Sequencing

PCR products were pooled and size-separated using a 1% agarose gel run at 145V. After visualization with SYBR™ Safe™ (Thermo Fisher) under blue light, DNA bands of around 450bp were cut out and purified using GeneJET gel extraction kit according to the manufacturer’s protocol (Thermo Fisher). DNA was eluted in water and then precipitated by adding 0.1 volumes of 3 M NaOAc (pH 5.2) and 1.1 volumes of isopropanol. After centrifugation for 10 min at 4°C, the resulting pellets were washed once in 70% EtOH and then air-dried. Afterwards a total of 100 μL water was added, the non-soluble fractions were spun down and removed, and the DNA concentration was quantified using a Qbit4 spectrophotometer system (Thermo Fisher). Libraries were quantified using VAHTS Library Quantification Kit (Vazyme) and deep sequencing was performed according to the manufacturer’s protocol using the MiSeq (Illumina) benchtop sequencing system. Data were obtained in FASTQ format and analysed with Outknocker (Schmid-Burgk et al. 2014).

### Immunofluorescence

For immunofluorescence staining, cells were fixed in 4% PFA-PBS for 10 min. After washing 3x with PBS, cells were blocked with 3%BSA diluted in PBS for 1h at RT and then incubated with anti-CD-19-PE antibody (1:50) in 3% BSA-PBS (Miltenyi Biotec) for 1h at RT. The nuclear stain Hoechst 33258 (2ug/mL, Sigma) in PBS was added for 10 min. Fluorescent images were obtained using a fluorescence microscope Leica DMi8 (Leica). To determine the number of CD19^+^ cells relative to the total number of cells, the total integrated density of the CD19 antibody was calculated and divided by the integrated intensity of the nuclei staining using ImageJ.

## Acknowledgements

Carina Gude, Marie Brauers, Olivia van Ray, Björn Hiller and Sara Desideri for technical assistance and help with the manuscript.

## Author contributions

SM and HR performed some of the experiments, optimized the technique and analyzed the data, ZK designed the work and interpreted the results, AR conceived and designed the work, optimized the technique, acquired data, interpreted the results and drafted the manuscript. SM, HR, ZK, JK and AR edited and revised the manuscript and approved the final version.

